# A Dynamical Systems Approach to Optimal Foraging

**DOI:** 10.1101/2024.01.20.576399

**Authors:** Siddharth Chaturvedi, Ahmed El-Gazzar, Marcel van Gerven

## Abstract

Foraging for resources in an environment is a fundamental activity that must be addressed by any biological agent. Modelling this phenomenon in simulations can enhance our understanding of the characteristics of natural intelligence. In this work, we present a novel approach to model foraging in-silico using a continuous coupled dynamical system. The dynamical system is composed of three differential equations, representing the position of the agent, the agent’s control policy, and the environmental resource dynamics. Crucially, the control policy is implemented as a parameterized differential equation which allows the control policy to adapt in order to solve the foraging task. Using this setup, we show that when these dynamics are coupled and the controller parameters are optimized to maximize the rate of reward collected, adaptive foraging emerges in the agent. We further show that the internal dynamics of the controller, as a surrogate brain model, closely resemble the dynamics of the evidence accumulation mechanism, which may be used by certain neurons of the dorsal anterior cingulate cortex region in non-human primates, for deciding when to migrate from one patch to another. We show that by modulating the resource growth rates of the environment, the emergent behaviour of the artificial agent agrees with the predictions of the optimal foraging theory. Finally, we demonstrate how the framework can be extended to stochastic and multi-agent settings.

**Author Summary:** Intelligence is a phenomenon that arises due to the interactions of an agent’s dynamics with the environment’s dynamics under the assumption that the agent seeks optimization of certain objective. Modelling both these dynamics as a single coupled dynamical system can shed light on patterns of intelligence that unfold in time. This report aims to provide a minimal in-silico framework that models the main components involved in natural phenomena, like optimal foraging, as a coupled dynamical system. Interestingly, we observe similarities between the surrogate brain dynamics of the artificial agent with the evidence accumulation mechanism that can be responsible for decision-making in certain non-human primates performing a similar foraging task. We also observe similarities between trends prescribed by theories prevalent in behavioural ecology such as the optimal foraging theory and those shown by the artificial agent. Such similarities can increase the predictability and explainability of artificial systems. We can now expect them to mimic these natural decision-making mechanisms by replicating such trends and we can thus understand the reasoning behind their actions. They can also increase the confidence of researchers to consider using such artificial agent models as simulation tools to make predictions and test hypotheses about aspects of natural intelligence.

## 1 Introduction

One way to define natural intelligence is by labelling it as the innate problem-solving abilities of biological agents under teleonomic assumptions [1]. It can either be studied by attempting to reverse engineer its functionality using (neurobehavioural) observational data in behaving biological agents or by designing and simulating adaptive artificial agents in complex environments to capture mechanisms of natural intelligence [2]. In this work, we are motivated by Braitenberg’s law of uphill analysis and downhill invention which states that it is easier to understand complex systems by reconstructing them from basic components rather than by estimating them from observations [3]. Thus, we seek to explore the latter option of modelling traits of natural intelligence from basic building blocks, as an approach to foster their understanding. We focus on one of the most prevalent traits of natural intelligence, that is, its ability to optimize the act of foraging for resources from its environment in order to satisfy the organism’s internal needs.

Foraging refers to the act of searching and gathering resources from the environment. It is ubiquitous and ancient in the sense that organisms ranging from unicellular organisms like bacteria [4, 5] to organisms having complex social structures like primates [6–8] display the urge to forage optimally. Various models depicting different aspects of foraging have been proposed under the umbrella of foraging theory [9]. Foraging models have uniquely predicted certain effects and phenomena regarding natural intelligence such as the effect of a forager’s internal energy budget on its sensitivity to the variance in reward or describing the environmental conditions under which a forager should or should not engage in exploration [9].

Patch foraging refers to one such model wherein a forager needs to decide when to quit foraging from a particular patch of resources and migrate to a different one. The caveat lies in the fact that often the value of resources in these patches is modeled as an exponentially decaying function of time [10]. This aspect of ceasing to consume the current resource for better prospects in a patch foraging task captures a large set of decision-making problems that natural agents face on a regular basis. Examples include, foraging for information instead of food or the act of quitting a problem-solving technique midway in order to explore a new one [11]. It has been shown that foraging activities employ multiple neural systems in the brain [12] to engage a variety of cognitive functions like learning of resource distributions across spatio-temporal scales and route planning [13].

Many optimal solutions and concrete behavioural characteristics of natural agents excelling in foraging tasks already exist and are studied extensively as a subset of optimal foraging theory [9, 14]. Particularly, in the case of patch foraging, the marginal-value theorem (MVT) dictates the optimal policy under some constraints and assumptions [10]. In brief, it states that an agent should leave a patch when the instantaneous rate of return from the patch falls below the long-term average rate of return of the environment. Many animals have been shown to comply with MVT in their natural settings [15–20]. However, there has been some push-back against MVT as it has been observed that temporal-discounting of the resource value promotes overstaying of an animal in the patch [21–23]. Furthermore, there exists some ambiguity in the value of the time horizon for which the average rate of reward of the environment is calculated [9]. In a trivial patch foraging task, it is assumed that the forager seeks to maximize the instantaneous rate at which it consumes resources without any other constraints being imposed [10]. Such models can easily be extended to more complex real-world scenarios. For instance, instead of following the trivial objective, a biological forager may try to maximize an additional objective to retrieve more resources to its place of retreat or nest. In such cases, the objective function of a traditional patch foraging task model can be augmented to include this extended constraint. Such models, where a place of retreat imposes constraints on the foraging activity, fall under central-place foraging tasks [24, 25]. Another frequently modelled constraint is the effect of multiple foragers foraging in the same environment, on a forager’s policy. The presence of other foragers in a patch can make it sub-optimal for a forager to visit it, even though the patch may be of higher quality than others. Further, given that each forager seeks to maximize its instantaneous rate of resource consumption, they should divide themselves among the patches as per the ideal free distribution theorem of the optimal foraging theory [26]. Thus, by augmenting components used to model a traditional patch foraging task, a similar model can also be developed for more complex scenarios where central-place foraging or the ideal free distribution can be observed [9].

Traditionally, patch foraging models have been designed and simulated using fixed decision rules and constraints, such as fixed travel-time between patches [10, 27, 28]. Work has also been done in formulating a normative theory of patch foraging decisions by posing it as a statistical inference problem [29]. More recent work has focused on using reinforcement learning (RL) to learn adaptive foraging behaviour [21, 30–34].

Hebbian learning has also been used to uncover an RL-like controller that displays adaptive foraging behaviour [35]. Work has also been done towards learning parameters of a general neural substrate based on Hebbian learning that forages optimally [36]. The patch foraging task can be considered as an example of a complex system [37] with its components being all the salient agent and patch behaviours needed for foraging to emerge. In this work, we propose to model a patch foraging task using a coupled dynamical systems approach which has extensively been applied to model complex systems like population persistence [38], celestial mechanics [39], climate modeling [40], coupled harmonic oscillators [41], and nonlinear electronics [42], to name a few. This approach breaks down the complex system into individual components, each with its own differential equation. These equations are then linked to mimic the system’s overall dynamics which further enables to gauge the effect of dynamics of other components on that of any individual component. It is important to note that other modelling techniques like agent-based models [43], cellular-automata [44] and network models [45] can also be used to model complex systems. They have also been used to model foraging tasks [46–48]. However, this particular approach allows the direct injection of known and informed model priors in the form of differential equations on each model component’s dynamics, making the model more ‘white-box’ in nature. Another motivation for using this approach can be derived from the fact that many studies of neurons recorded in the cortex reveal the existence of complex temporal dynamics in their firing [49], and these dynamics are heavily influenced by the dynamics of the environment in which they are embodied [50, 51].

We divide the complex system into three components, namely the brain dynamics of the agent (controller), the position dynamics of the agent, and the growth dynamics of the resources. Many modelling techniques to formalize an agent’s control policy, ranging from linear proportional-integral-derivative (PID) controllers [52, 53] to more general recurrent neural networks [54, 55] exist. Here we opt to use a neural differential equation (NDE) [56] to model the agent’s brain dynamics. This choice is convenient from the neuroscientific standpoint as it allows the use of a brain-inspired differential equations such as rate-based models that capture average neuronal population activity [57] as a prior on brain dynamics. Interpreting the brain of the agent as a dynamical system whose states evolve across time is the topic of dynamical systems theory in neuroscience [58, 59]. Additionally, it allows the use of deep neural networks (DNNs) as function approximators to estimate the differential term which, in turn, provides high capacity function approximation to an arbitrary level of precision that can be optimized using automatic differentiation [60]. Here we use a modern interpretation of a Wilson-Cowan model [49], traditionally used in neuroscience to model firing rates of neural populations. We allow the parameters of this NDE to be adjustable so that they can be estimated by optimizing for the optimal foraging objective of maximizing the rate of resource consumption while minimizing the energy expenditure.

The dynamics of the resources growing in the environment are modelled using a Lotka-Volterra (LV) model [61]. The LV model is parameterized by a growth rate and a decay rate which together dictate the carrying capacity of the population. Finally, the agent’s locomotion in the environment is defined by a double-integrator (DI) model, describing the equations of motion of a particle in two-dimensional space [62].

In summary, through this work we introduce the modelling approach based on posing a complex system as a coupled dynamical system, to the task of patch foraging by artificial agents. Crucially, we model the artificial agent’s brain dynamics based on a biologically inspired NDE to capture the similarities between real and simulated neurobehavioral data. This is reminiscent of neuronal dynamics observed in single-cell recordings from non-human primates while patch-foraging in a computerised task [63].

We also observe a shift in patch-residing times of the agent when the growth rate of the resources in the environment is modulated. This observation aligns with the predictions of the optimal foraging theory. That is, a forager tends to overstay in a patch when the environment becomes poorer in terms of resources [9, 10]. Further, we extend the modelling approach to multi-agent and stochastic settings to show how it can be augmented to accommodate modeling of more complex scenarios.

This paper is structured as follows. Section 2 describes the formulation of the coupled dynamical system and the process used for its optimization. Section 3 presents all the experiments performed using this setup. Finally, Section 4 discusses empirical findings and lays down directions for future work.

## 2 Methods

### 2.1 Modelling the dynamics

Our setup is a coupled dynamical system consisting of three components: A position model, a control model, and a resource model. The position and control model together comprise the agent whereas the resource model comprises the environment. Each component is modeled as a dynamical system, and their interactions are captured by a set of coupled differential equations. Specifically, in our approach, we model everything in terms of forced ordinary differential equations (ODEs) of the form

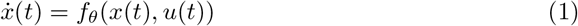

where *x*(*t*) is the state of the system at time *t*, 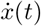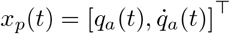 is its time derivative, *u*(*t*) is the forcing function, *f* is the state equation that models the system dynamics and *θ* are (optional) free parameters. For conciseness, we often omit the time index from our notation. Based on this general notation, we specify the different model components.

#### 2.1.1 Position model

The agent is modelled as a particle moving on the surface of a torus. The toroidal environment is chosen to remove spatial boundary conditions and facilitate a continuous state space for the agent’s position. This serves as a basic representation of an agent moving in 2D space. An overview of the setup can be seen in Fig 1(a).

**Fig 1.**
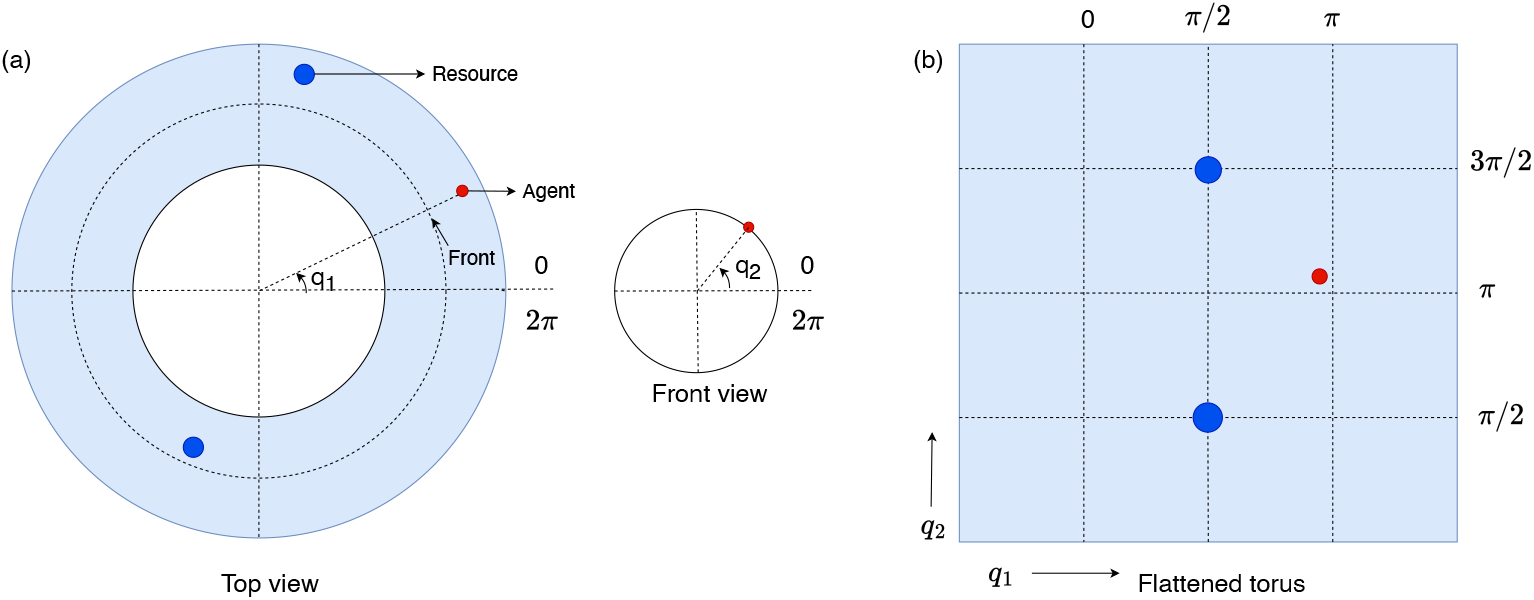
A toroidal environment with two degrees of freedom. (a) The top view and the front view of the environment. The agent is depicted with a red disc and the resources with blue discs. The resources are growing at random locations. (b) A flattened view of the torus with the resources now growing at (*π/*2, *π/*2) and at (*π/*2, 3*π/*2).

The motion of the agent is modelled using a deterministic double integrator [62], which describes how its position, velocity and acceleration are related. This modelling technique is prevalent in many real-world scenarios like automotive and robot control where the control input directly influences the acceleration of the system [64]. It implies that the dynamics of the position model *f*_*p*_ is given by

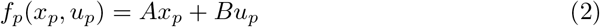

where 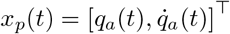 with *q*_*a*_(*t*) *∈* ℝ ^2^ being the position of the agent in 2D space. Here we employ the common convention of including both the position and the velocity in the state space of the model to accurately track the position of the forager and convert the second-order dynamics to first-order dynamics [65]. The system matrices are given by

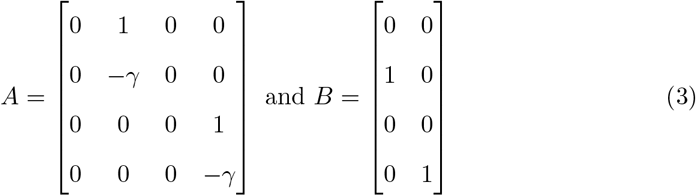

where *γ* is a damping factor. *A* and *B* represent the state matrix and the input matrix of the model respectively, which are used in a typical state-space model. We further assume that the control vector *u*_*p*_(*t*) *∈* ℝ ^2^ is a readout of the neuronal firing rates *r*(*t*)given by

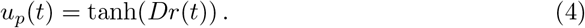

Here, *D ∈* ℝ ^2*×n*^ is a readout matrix, mapping the firing rates of *n* neurons to the accelerations. The tanh function ensures that the magnitudes of the accelerations along both the directions remain bounded. Firing rate dynamics are described in Section 2.1.3.

#### 2.1.2 Resource model

We model the distribution of resources by choosing *m* = 2 fixed locations on the surface of torus which are capable of growing resources as shown in Fig 1(b). These locations are given by (*π/*2, *π/*2) and (*π/*2, 3*π/*2). The information about the location of the resources is stored in a constant vector *q*_*r*_ *∈* ℝ ^4^. Thus, in this model the dynamics only exist in terms of the value of these resources *x*_*r*_(*t*) = [*s*_1_(*t*), *s*_2_(*t*)]^*⊤*^, where *s*_*i*_(*t*) *∈* R is the value of the *i*-th resource. Further, these dynamics are based on the Lotka-Volterra (LV) model [61], wherein the value of resources keeps growing autonomously to a maximum carrying capacity. These values are also depleted in direct proportion of the distance between the resource and the forager. The resource value dynamics *f*_*r*_ are given by

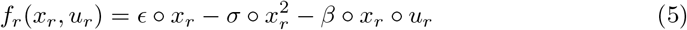

where *ϵ ∈* ℝ ^+^ is the constant growth rate, *σ ∈* ℝ ^+^ models the constant decay rate and *β ∈* ℝ ^+^ models the constant consumption rate of the resources. The operator *○* represents an element-wise multiplication. The control vector *u*_*r*_(*t*) = *w*(*q*_*a*_(*t*), *q*_*r*_), *∈* ℝ ^2^ represents the distance between the agent’s position *q*_*a*_(*t*) and the (fixed) resource locations *q*_*r*_. The distance function *w* is computed based on the sines and cosines of *q*_*a*_(*t*) and *q*_*r*_ so that the continuous nature of the agent’s position with respect to the resources remains intact and boundary conditions are not imposed. The structure of *w* is detailed in Eq 15 of the S1 file.

#### 2.1.3 Control model

The controller model represents a surrogate brain model which determines how the agent controls its position based on its observations. The controller dynamics *f*_*c*_ are modeled using a modern interpretation of a Wilson-Cowan model [49] which is a cornerstone in computational neuroscience for modelling the dynamics of neuronal populations [66]. It provides a basic framework for simulating the complex dynamics of neuronal populations, capturing the aggregate interactions and oscillatory behavior essential for understanding brain function and its emergent properties. The controller dynamics are given by

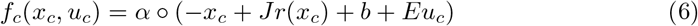

where the system’s state *x*_*c*_(*t*) = [*z*_1_(*t*), *z*_2_(*t*), …, *z*_*n*_(*t*)]^*⊤*^ *∈* ℝ ^*n*^ represents the summed and filtered synaptic currents *z*_*i*_ of *n* neurons and *r*(*x*_*c*_(*t*)) = tanh(*x*_*c*_(*t*)) represents their firing rates. The firing rates are also used as the control input to the position model in Eq 4. The recurrent weights of the controller dynamics are given by *J ∈* ℝ ^*n×n*^, *b ∈* ℝ ^*n*^ is a bias term and *E ∈* ℝ ^*n×p*^ are the input weights. The controller dynamics are further determined by 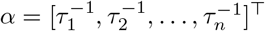, where *τ*_*i*_ *∈* ℝ ^+^ is the time constant for the *i*^*th*^ neuron. Importantly, the control model can be inferred as a neural differential equation (NDE) with *J* representing the inter-neuron connections and *E* representing its interface with the observations.

In this case, the control vector *u*_*c*_ *∈* ℝ ^*p*^ is a vector of *p* observations sensed by the agent. These observations inform the control model about the system’s current state which is in turn used by it to produce the next action. They consist of the sines and cosines of the agent’s position, the velocity of the agent in the 2D space, the value of the resources, and the net amount of resources consumed by the agent at time *t* as detailed in the following section. Its exact form is described in Eq 18 of the S1 file. The initial value of the state vector is obtained by passing the control through a linear readout matrix *R* such that *x*_*c*_(0) = *Ru*_*c*_(0). Further, all the learnable parameters of the position and control models form a subset of the learnable parameters *θ* such that *θ* = *{D, J, b, E, α, R}*. An overview of how the three models are inter-connected together to formalize the coupled dynamical system is illustrated in Fig 2.

**Fig 2.**
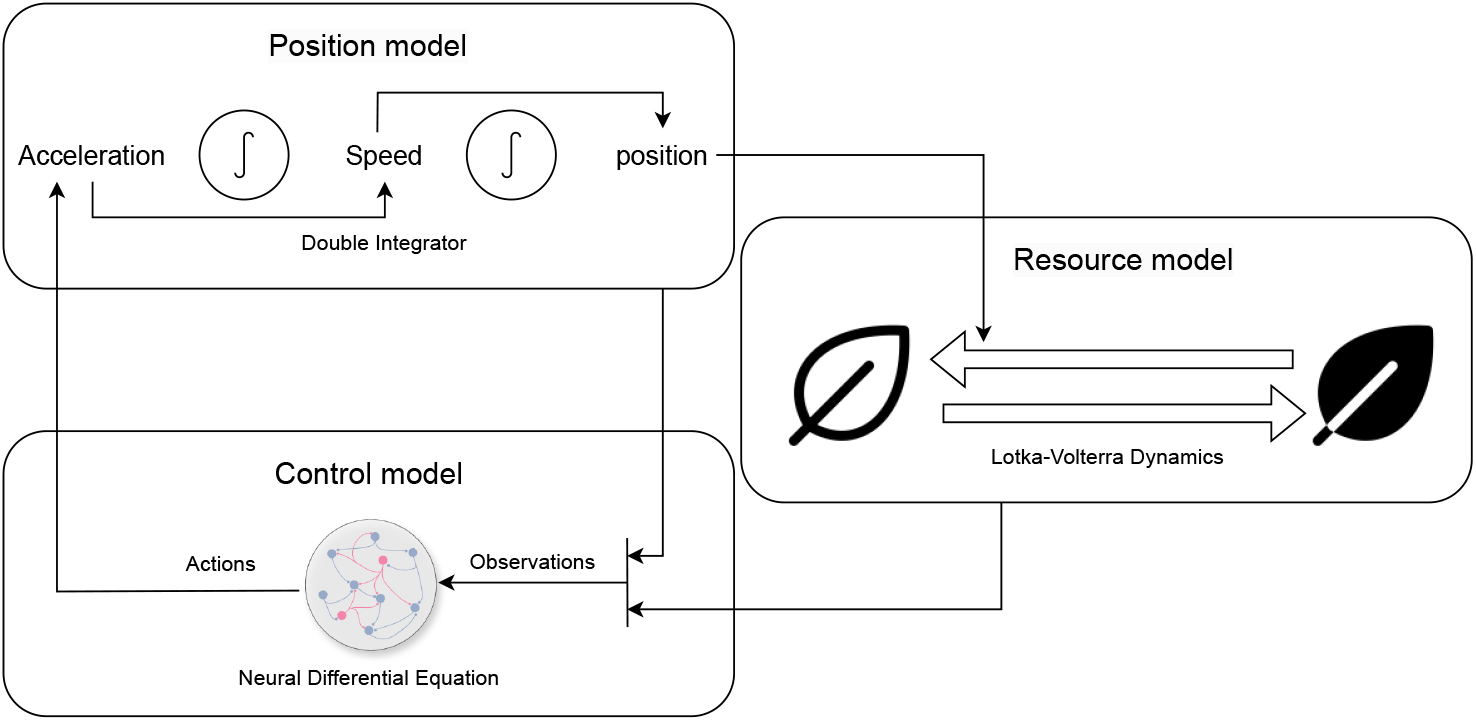
Block diagram showing all the components connected together.

#### 2.1.4 Stochastic dynamics

To implement stochastic dynamics in the system we augment Eq 1 in the context of the resource model (Section 2.1.2), to include a noise term, making the equation a stochastic differential equation (SDE) such that

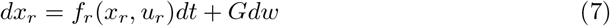

where the vector *x*_*r*_ represents the values of the resources, *f*_*r*_ represents the drift term of the resource model dynamics, *G* is a constant matrix which determines how the noise effects the dynamics of the model and *w ∈* ℝ ^2^ is a multivariate Wiener process [67].

Thus, *x*_*r*_ which is a subset of the observation vector used by the control model in Section 2.1.3, now includes stochasticity. This makes the agent observations noisy. Following such a methodology of appending a noise term after the drift term, the deterministic dynamics of the control model and the position model can also be augmented to account for stochasticity.

### 2.2 Optimal foraging objective

We aim to optimize the agent in order to evolve it into a rate-maximizing forager. Thus, the aim of the agent is to maximize the average amount of resources it consumes over time and minimize the amount of control exerted. That is, the goal is to learn a policy parameterized by *θ* such that

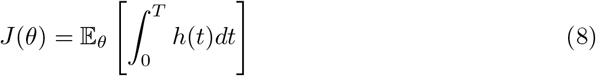

with 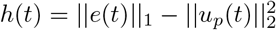 is maximized. Here, *e*(*t*) = *β ○ x*_*r*_(*t*) *○ u*_*r*_(*t*), represents resource consumption, as presented in Eq 5. We also impose a cost on movement by penalizing the square of control input *u*_*p*_ of the position model in Eq 2. Finally, the problem statement becomes finding the optimal value of *θ*, labelled as *θ*^*∗*^, such that.

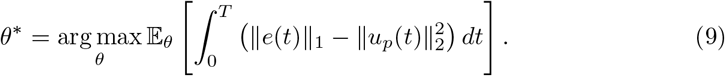

### 2.3 Multi-agent system

In this section we show how can we extend the modelling approach to account for a multi-agent system with *K* agents.

#### 2.3.1 Position model

The position model as depicted in Section 2.1.1 can be extended to a multi-agent setting by introducing the position and velocity of additional agents as a different vector of state-space variables. Thus the position model based on Eq 2 transitions to

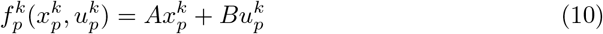

where *k ∈ {*0, *K}* represents the position model for the *k*^*th*^ agent. The acceleration of each agent is given by 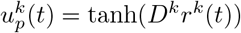 wherein *D*^*k*^ represents the readout matrix which is also included in the set of learnable parameters *θ* of the model.

#### 2.3.2 Resource model

In a multi-agent setting the resource model in Section 2.1.2 is extended by accounting for the resources consumed by all the *K* agents active in the environment. Thus the resource model presented in Eq 5 becomes

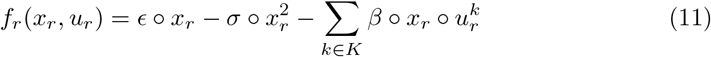

where 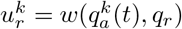 represents the distance function for *k*^*th*^ agent as detailed in Eq 15 of the S1 file.

#### 2.3.3 Control model

Similar to the position model, the control model for a multi-agent setting can be formulated by simulating a control model for each active agent. In particular, in the case of the control model, this entails including the learnable parameters of each agent in the set of learnable model parameters *θ*. Thus, the control model as presented in Section 2.1.3 becomes

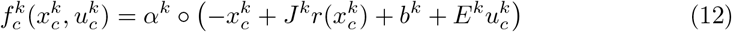

for the *k*^*th*^ active agent. Further, each *α*^*k*^, *J*^*k*^, *b*^*k*^, *E*^*k*^ also becomes part of the learnable parameters *θ*.

#### 2.3.4 Optimal foraging objective

We augment the objective function as introduces in Section 2.2 by simply summing over a similar foraging objective for all the *K* agents, thus the objective function becomes

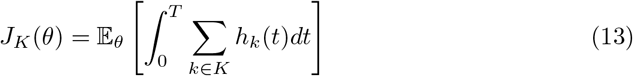

wherein we optimize for the optimal adjustable model parameter set *θ*^*∗*^ as

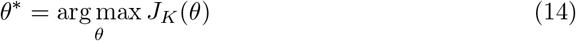

### 2.4 Learning procedure

Having access to the deterministic dynamics of the system, we use a simple learning algorithm based on the truncated backpropagation-through-time (t-BPTT) procedure [68] to find the optimal values of *θ*. Firstly, for each learning trial, we sample 64 initial points of the coupled dynamical system (see S1 file for details on system initialization). Following that we simulate the trajectories corresponding to the sampled initial points. Each trajectory is composed of 1000 time steps wherein the initial step size is set to 0.04*s*. Thus, each trajectory represents the simulation of the system for 40*s*. For computing the gradients according to the t-BPTT algorithm, the truncated interval is set to 200 time steps per trajectory. The value of *θ* is updated after every learning trial by averaging the gradients collected over all the trajectories sampled in that learning trial with respect to the objective function defined in Eq 8. The learning algorithm is depicted in Algorithm 1 and the details of the simulations can be found in the S1 file.

#### Algorithm 1

Learning Procedure

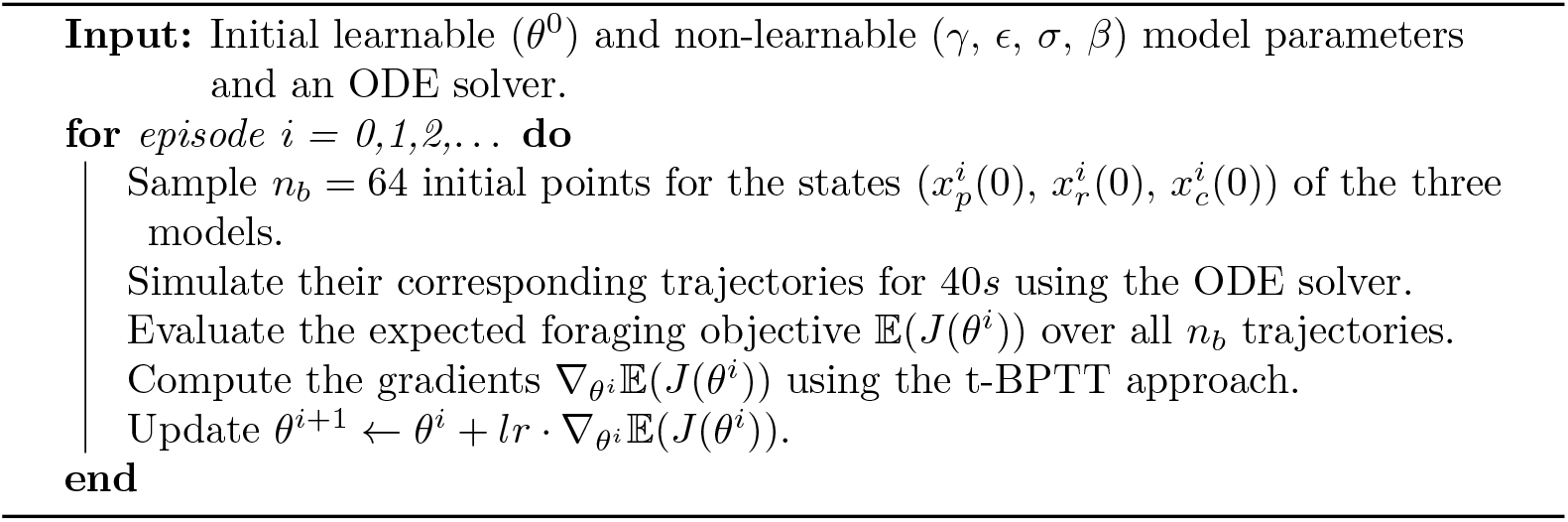

## 3 Results

### 3.1 Emergence of foraging

After optimizing the model’s learnable parameters *θ*, the optimal policy reflects the following behaviour qualitatively. Firstly, after being spawned at a random location, the agent migrates to a patch where a resource grows. We classify this behaviour as the initial migration. Then, the agent demonstrates position control around the patch and depletes the resource value in it to a particular level. We term this behaviour as position control. Finally, without completely depleting the resource value, the agent moves on towards the other patch. The agent then keeps repeating this behaviour by oscillating between the two patches. We label this behaviour as oscillatory migrations.

Fig 3(a) represents the policy consisting of the three constituent behaviours. Fig 3(b) shows the increase in the foraging objective while training, demonstrating effective learning of the task. Fig 3(c) shows behaviour of the agent at various stages of training, starting from a random agent to an optimized agent. The degree of optimization is reflected in the amount of resources gathered by the agent at various stages of training. It can be seen that the agent learns the act of migration between resource patches rather quickly. We observe that the bulk of training time is used up to optimize the patch residing time. Other than the amount of resources collected by the agent we note two more performance indicators as function of the training process. Firstly, the average amount of time taken for an agent to migrate towards one of the resources for the first time, after being spawned. This average is calculated by spawning the agent at five random locations selected in the beginning of the training phase. In our context, a visit to a resource is noted as a migration only when the agent stays in the close vicinity for at-least five consecutive steps. Secondly, the average mean squared error observed when the agent is performing position control near the resource. The agent is considered to be performing position control only if stays in the vicinity of 0.5 rad near either of the resource for at least ten consecutive time steps. Further, the average is calculated for each patch visit during a trial and for five different initial spawning position of the agent. Fig 3(d) shows both these performance indicators decreasing with training which logically aligns with the increase in the foraging objective.

**Fig 3.**
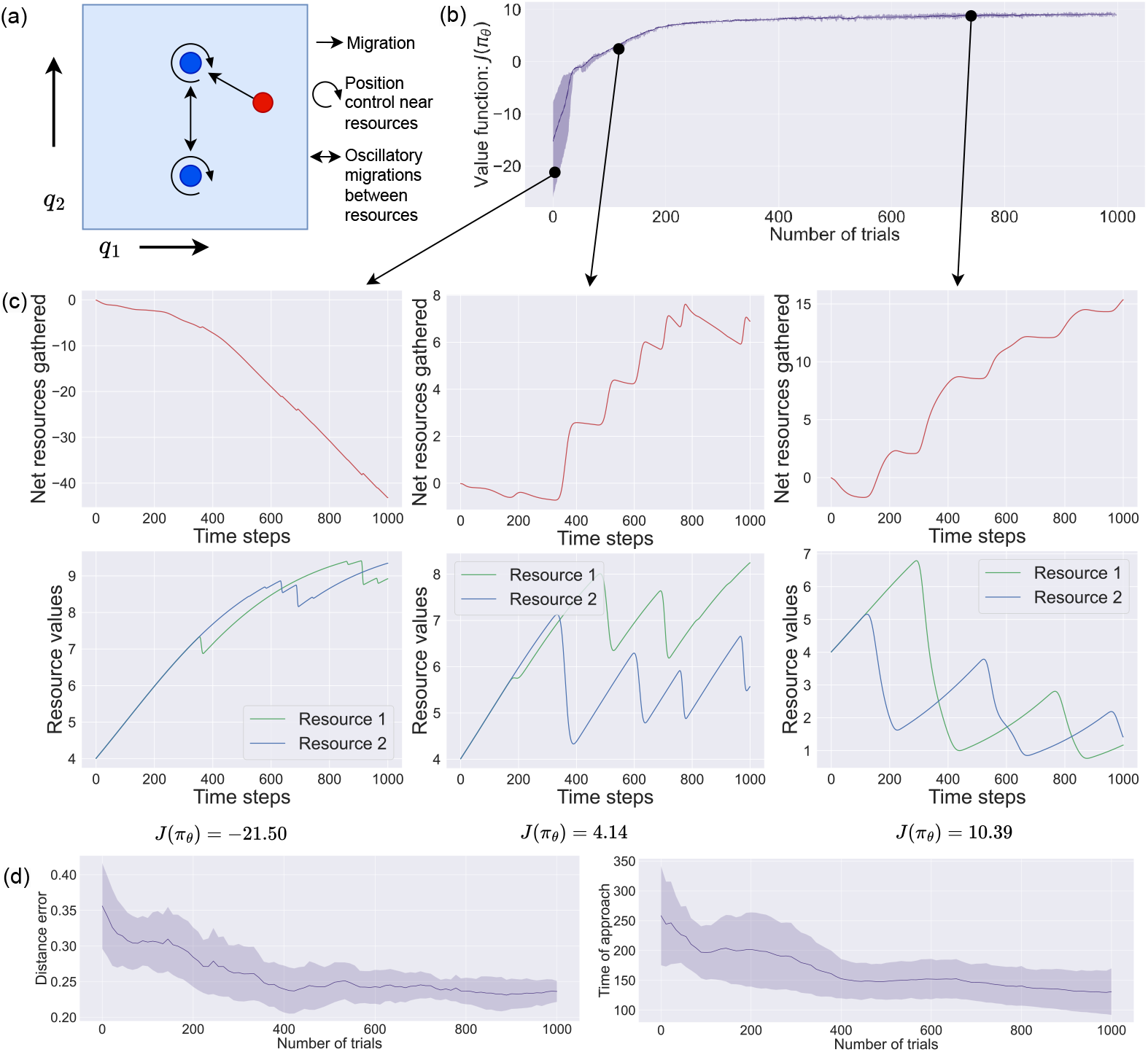
Results of learning. (a) Learned adaptive foraging behaviour. (b) Loss curve representing the objective function in Eq 8. (c) Net amount of resources collected by the agent and the resource values at various stages of training for a sampled trial. (d) average time of the first approach and average position error as a function of the training process.

Since in the setup, the dynamics of the resource values in the future depends on the agent’s current patch residing time, the calculation of optimal patch residing time using MVT is not a straightforward task [9]. The reason being that overstaying in a patch can translate to more time for the other patch to regenerate. On the other hand, staying in a patch for too long can lead to the loss of opportunity of gaining more resource value at a faster rate. This trade-off may lead to a general overstaying of the agent in a patch with respect to the optimal time of leaving as predicted by the MVT. As seen in Fig 3(c), in general, for an optimal agent, the patch residing times increase with each patch visit as the environment becomes poorer in terms of the resource values.

### 3.2 Evidence accumulation

Neural recordings obtained from the dorsal anterior cingulate cortex (dACC) of non-human primates that were tasked to carry out a computerized patch foraging task, suggest that the underlying computations used by some neurons in the area, for deciding when to leave a patch, may represent an evidence accumulation mechanism [63]. This mechanism generally employs the use of a decision variable, where the value of the variable represents the amount of evidence in support of a particular decision for instance when to leave a patch. This value accumulates over time and once it reaches a certain threshold, the decision is made. A decision is made sooner if the rate or slope at which evidence is accumulated for it is in general greater [69, 70]. This is shown in Fig 4(a), wherein the red red trajectory reaches the threshold value in the least amount of time, thus a decision in this case will be taken the soonest. Moreover, a decision can also be made sooner if the amount of evidence required for it is lesser, in other words, the magnitude of the threshold that needs to be crossed by the decision variable is lesser [71] as shown in Fig 4(b).

**Fig 4.**
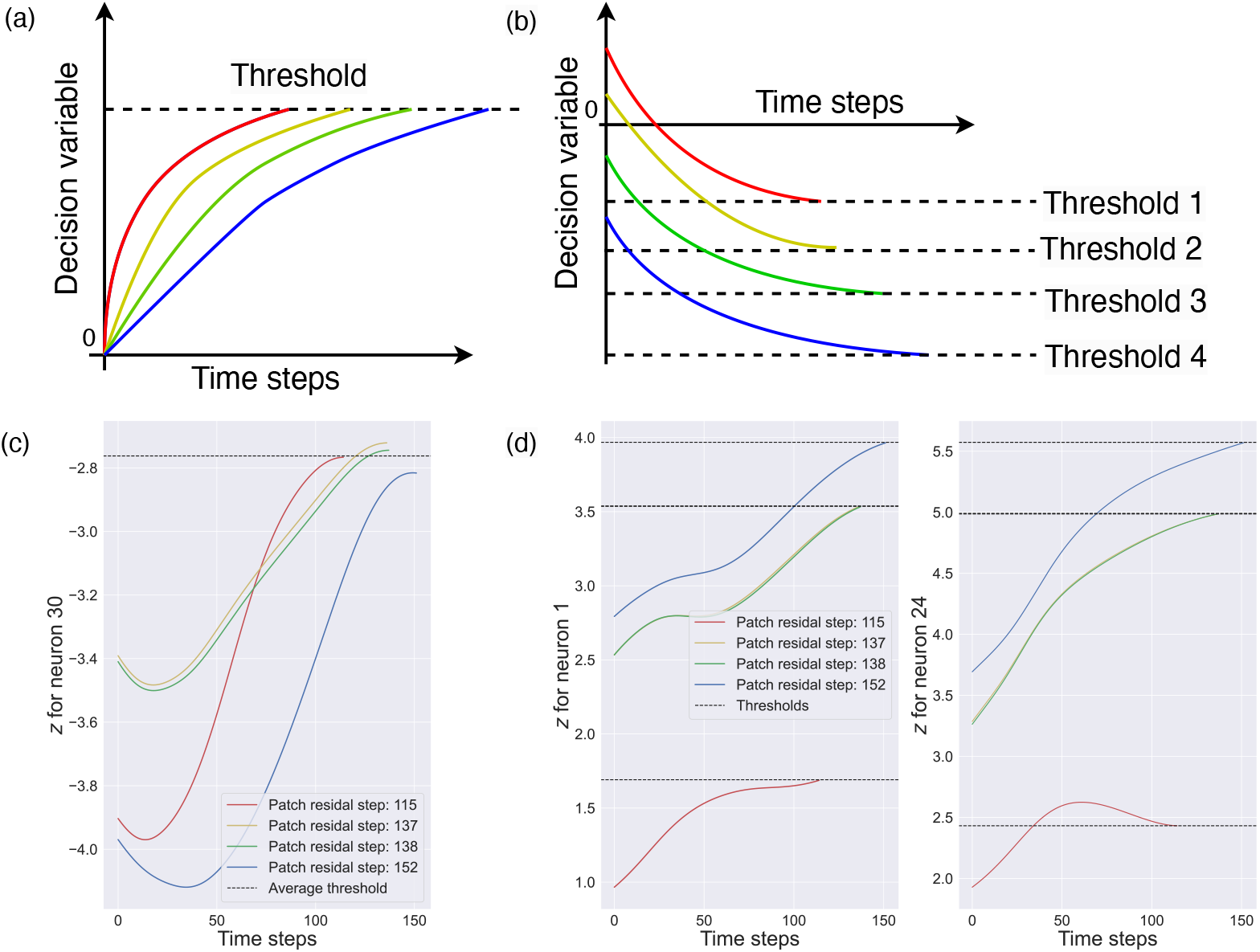
Evidence accumulation. Evidence accumulation mechanism in an ideal case: (a) Higher slope (b) Higher threshold magnitude. (c) and (d) Value of the neuron state *z* in Eq 6 for three empirically selected neurons for four sampled trajectories having different initial conditions. The trajectories are cropped from the times of second patch entry to the times of second patch departure. In (c), we observe faster evidence accumulation via higher slope value i.e. the trajectory displaying the highest slope value reaches the Average threshold value and exits the patch the soonest. In (d) we see the other version of evidence accumulation where the magnitude of the threshold value for the earlier patch departure is set to a lower value.

As Fig 4(b) shows, in this case, the threshold beyond which a decision is taken, is itself set at a lower magnitude for the case where the decision is taken sooner. For instance, the threshold value for the red trajectory (Threshold 1) is set at the lowest magnitude, thus a decision for it will be taken the soonest. On the contrary, in the case of the blue trajectory the magnitude of the threshold value (Threshold 4) is the highest, thus the decision in this case will take the most time. Evidence accumulation has been nominated as a general decision-making mechanism that underlies many value-based decisions in organisms including patch foraging decisions of leaving a patch [72]. This mechanism is particularly expected to emerge in adaptive agents when the environment is partially observable [73]. This aligns with the design of our setup, as in our case the agent can observe the values of the resources but not their positions which might lead to the need of evidence accumulation for the decision of when to leave a patch or resource. We observe a similar trend in neurons of an optimized agent in the setup.

Fig 4(c) and (d) show the dynamics of some empirically selected neurons from the control model described in Section 2.1.3, for different sampled trajectories during the second patch visit. In Fig 4(c) it can be seen that the neuron accumulated evidence with a higher slope when the patch residing time is lesser. In contrast, in Fig 4(d) we observe the another evidence accumulation mechanism wherein the value of the threshold beyond which the agent decides to leave the patch, is set at a higher magnitude for longer patch residing times. Further, when the patch residing time is very similar (137 and 138 for the yellow and green trajectories respectively in Fig 4(c) and (d)), the neuronal dynamics follow a very similar curve while respecting the trends used by the evidence accumulation mechanism to undertake a decision. This suggests that the learned control model uses a mechanism very similar to evidence accumulation.

### 3.3 Different growth rates

A salient conclusion of optimal foraging theory, regarding patch foraging, is that the optimal time of residing in a patch for a forager increases if the environment becomes poorer in terms of resources [9, 10]. The environment can become poor in resources if the distance between the patches or the travel time between them grows. Alternatively, if the maximum possible value of the resources in the patch becomes less, the environment can be termed poorer in terms of resources. In this experiment, we trained agents to forage in three different environments with different growth rates of the resources in the patches. This allowed us to see if the simulation results align with the OFT predictions. Environments share a common value of the decay rate while using different growth rates. This translates into different carrying capacities of the patches in each environment.

Fig 5(a) shows a comparison of the average patch residing times calculated for the first three patch visits across different trajectories in the three environments (mean of 143, 107.33, and 101.4 time-steps for growth rates of 0.8, 1.0, and 1.2 respectively).

**Fig 5.**
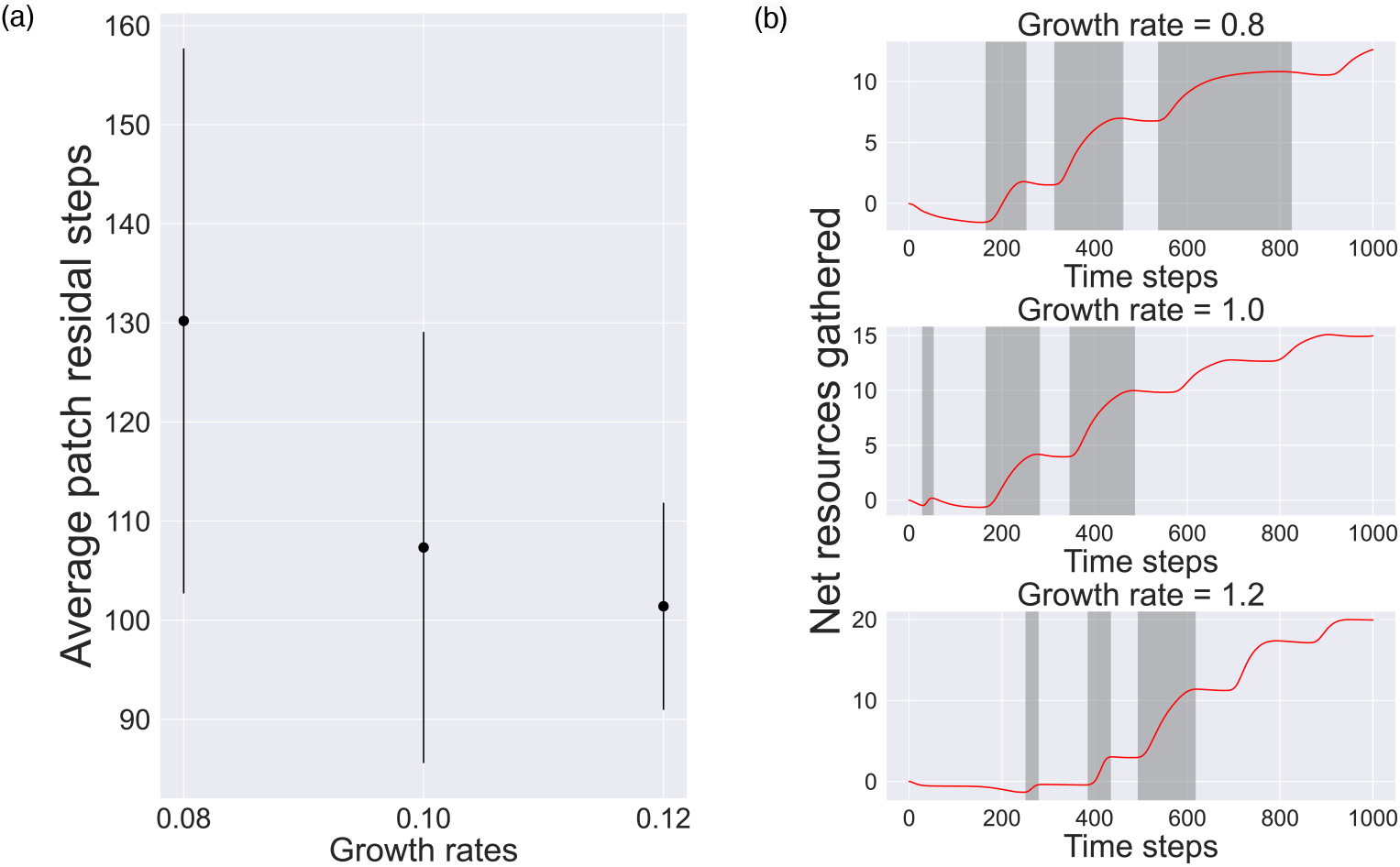
Comparisons of foraging behaviour in three different environments. (a) Average patch residing times for the first three patch visits for different trajectories sampled in the three environments. (b) Net amount of resources gathered by the agents during a sampled trial in each environment. The grey regions indicate the first three patch residing periods, which decreases for the sampled trial as the growth rate increases across environments (mean of 173.0, 92.33 and 65.67 time-steps for growth rate of 0.8, 1.0 and 1.2 respectively).

Results show that the mean patch residing time for the agent decreases as the environment becomes richer in resources. In Figs 5(b), it can be seen that the environment with higher growth rate generates more resources, which, in turn, leads to an optimal agent collecting more resources and spending less time in a patch.

### 3.4 Stochastic observations

To model a setup wherein the agent receives noisy information about the resource values we replace the deterministic resource model with Eq 7. For training this stochastic system we use the same procedure as described in Section 2.4, but we substitute the Tsit5 solver with an EulerHeun solver [74]. This allows us to account for the stochasticity of the system.

In Fig 6(a) we train a model with *G* = 0.1*I* where *I* is a two-dimensional identity matrix, and compare the displayed position error as introduced in Section 3.1 with that of a completely deterministic system. This performance indicator appears similar for both setups. We then compare the net amount of resources gathered by an agent optimized in an environment which produces stochastic observations with that where the observations are deterministic. The former does better than the latter on average when tested in an environment with stochastic observations, as shown in Fig 6(b).

**Fig 6.**
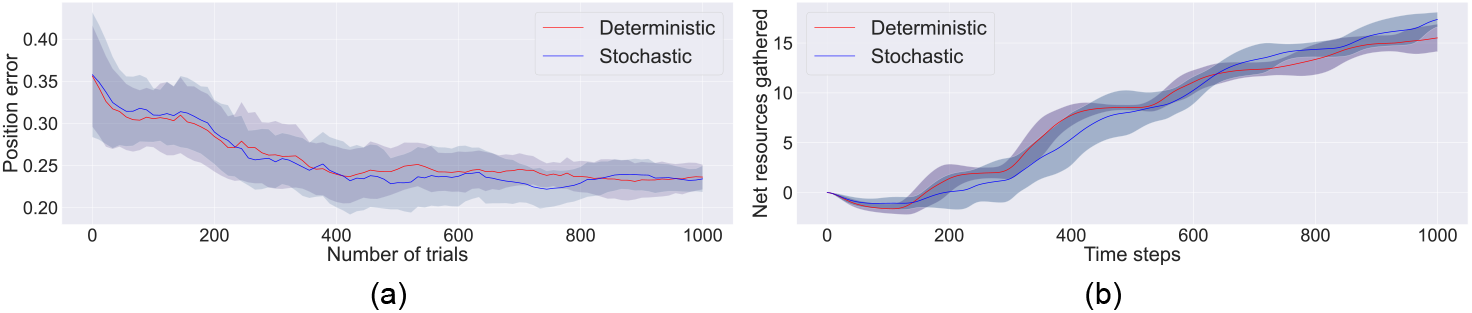
Performance comparison of a deterministic system and a stochastic system. (a) Position error during the training phase. (b) Foraging objective during a trial in a stochastic environment.

### 3.5 Multi-agent setup

In this experiment we show how foraging agents alter their choice of foraging patch based on different growth rates of the resources when they forage in presence of other agents. Here, we employ the methodology described in Section 2.3 to model a multi-agent setup for *K* = 2 agents and optimize it for three different environment settings pertaining to the growth rates of the resources. Namely, when the growth rate of one of the resources at (*π/*2, *π/*2) remains constant at 0.1 and that of the other resource at (*π/*2, 3*π/*2) is varied across different values.

As shown in Fig 7(a), when the growth rates of both the resources are set equal at 0.1 the agents choose to migrate towards different resources aligning with the ideal free distribution theorem of optimal foraging theory. When the growth rate of the second resource (set at 1.0) is much higher than the first resource (set at 0.1) both the agents migrate towards it, as depicted in 7(b). Finally, when the growth rates are comparable, as in the case of 7(c), the agents display a more complex migration strategy. This complexity may be attributed to the dynamics embedded in the resource model. The interplay between the growth rate of resources and emergence of such complex behaviour can be an interesting topic of future investigations.

**Fig 7.**
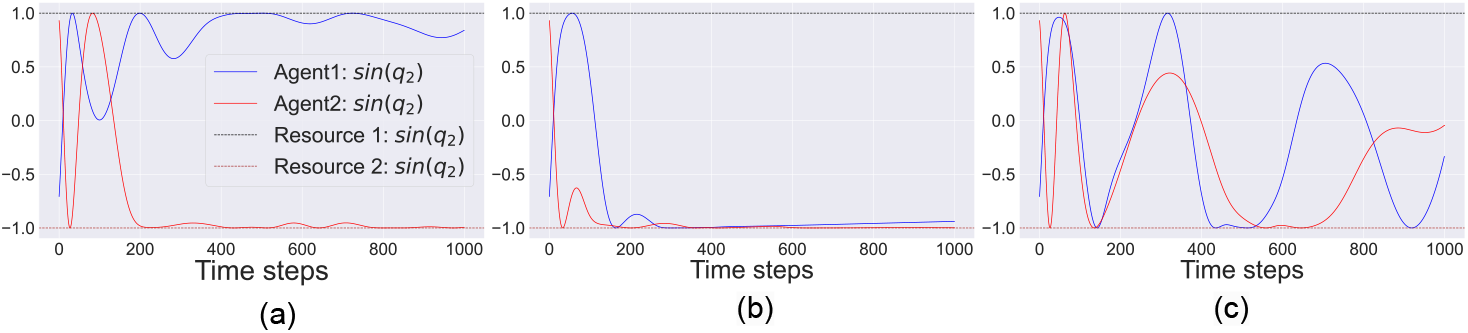
Comparison of optimized multi-agent policy with unequal growth rate of resources for a sampled trajectory, whereby resource 1 at (*π/*2, *π/*2) has a constant growth rate of 0.1 and the growth rate of resource 2 at (*π/*2, 3*π/*2) varies as (a) 0.1, (b) 1.0 and, (c) 0.3. The plots just show the *sine* of the positions of the resources and the agents along the *q*_2_ direction.

## 4 Discussion

We presented a general and minimal setup to model and simulate the dynamics involved when an adaptive agent learns to forage. The setup is general in the sense that we combine the salient aspects of the environment and agent dynamics that implement foraging at large, as one coupled dynamical system. This was done such that the continuous nature of the dynamics are preserved. The setup is minimal in the sense that, since we have the access to the complete deterministic dynamics of the system, simple supervised learning algorithms such as t-BPTT can be used to optimize the free parameters of the coupled system.

We showed that the optimized controller model, as a surrogate brain model, consists of neurons which display evidence accumulation dynamics that are similar to the dynamics observed in the dACC region of non-human primates that have learned to patch forage. Further, when the environment parameters like the growth rate of resources were tweaked, adaptive foraging behaviour matched the predictions made by optimal foraging theory. Namely, the agent stayed in the patches for longer periods of time when the environment became poorer in terms of resources.

We also demonstrated how this modelling approach can be extended to model more complex scenarios such as accounting for stochastic environments and multi-agent setups. In the context of a stochastic environment, our aim was to show an example of a setup where the agent receives stochastic observations. By augmenting different components of the model in a similar way one can account for stochasticity in terms of learned policy and agent actions. In the case of a multi-agent setting, we presented an example for a two agent setup, but by following a similar model augmenting technique, one can account for a setup with more number of agents. It is also to be noted that, as observed in the case of varying the growth rate of the resources, the qualitative nature of foraging policies show interesting transitions. These transitions can be attributed to the continuous nature of the resource dynamics and can be the scope of future studies.

Incorporating biologically-inspired brain models into artificial agents may shift them towards transparency and predictability because now we can expect similar trends in the process of their decision-making as their biological counterparts. In general, such trends allow us to use prior knowledge and our understanding of the mimicked natural models to anticipate the artificial agent’s behaviour and discern the reasoning behind its actions. Such transparency when combined with other white-box modeling approaches, for instance, passing control torques in robotics [75] can foster greater trust among its practitioners [76]. Thus, in this work, we hope to contribute towards the confidence of researchers to employ such biologically inspired artificial agents as simulation tools to study and test hypotheses about aspects of natural intelligence.

The advantage of our dynamical systems approach to optimal foraging is that complex emergent behaviour can be studied using a minimal setup which only requires the numerical simulation of a system of differential equations and the optimisation of free parameters using automatic differentiation. This setup facilitates hypothesis testing since assumptions about the agent and the environment translate into specific changes in the differential equations that make up the optimal foraging model.

The model presented, contains the minimal components needed for optimal foraging to emerge and lacks some of the realism that exist in nature. For instance, currently the agent receives complete and deterministic information about the environment. It will be interesting to study the change in dynamics of the agent’s brain when its sensors become noisy or when it receives partial observations from the world as in the case of natural organisms [77]. We may also extend the environment in various ways. Noisy resource dynamics can be facilitated by resorting to stochastic differential equations [29]. Modelling the distribution of resources across space can be handled using partial differential equations [78]. Downsides of these extensions are the increased computational complexity involved in simulating such systems.

Another interesting future avenue would be to extend the current setup to a multi-agent setup and analyse the multi-agent strategies that agents learn in order to forage together [79]. More specifically, the analysis of competitive and collaborative nature of these strategies may uncover some interesting parallels that are also observed in nature [80, 81].

Finally, the current setup estimates free parameters using automatic differentiation. Ultimately, gradient-free approaches such as evolutionary algorithms may prove to be more convenient as they do not require differentiation through the environment. Furthermore, this would allow studying how the system of differential equations evolves structurally under environmental pressure [82].

In conclusion, with the development of a minimal model of optimal foraging, cast in terms of a system of coupled differential equations, we aim to provide researchers with a new approach to studying how complexity emerges as natural agents adapt to their environment.

## Acknowledgments

This publication is part of the project Dutch Brain Interface Initiative (DBI^2^) with project number 024.005.022 of the research programme Gravitation which is (partly) financed by the Dutch Research Council (NWO).

## Supporting Information

**S1 File. Modelling details.**
The contents of this file represents some of the finer modelling details and empirical choices used to model the system.

