## Supporting Information for "A Dynamical Systems Approach to Optimal Foraging"

This section presents some of the finer modelling details and empirical choices used to model the system.

### Distance function

The function $w(q_a, q_r) \in \mathbb{R}^2$ used to compute the distance metric between the agent's position $q_a$ (at some time $t$) and the resource locations $q_r$ in Section 2.1.2 of the main text is given by

$$w(q_a, q_r) = \left[ e^{\frac{-l}{d(q_{r_1}, q_a)}},\ e^{\frac{-l}{d(q_{r_2}, q_a)}} \right]^\top \tag{15}$$

where $l$ represents the steepness of the Gaussian kernel and is set to $-7.0$ and $d(q_{r_i}, q_a) \in \mathbb{R}^+$ is given by

$$d(q_{r_i}, q_a) = ||\sin(q^1_{r_i}) - \sin(q^1_a),\ \cos(q^1_{r_i}) - \cos(q^1_a),\ \sin(q^2_{r_i}) - \sin(q^2_a),\ \cos(q^2_{r_i}) - \cos(q^2_a)||_2\,. \tag{16}$$

Here, $q_{r_i} = (q^1_{r_i}, q^2_{r_i})$ represents the coordinates of the resources $r_i$ along the two axes of the torus. For the two resources growing in the environment these are given by $q_{r_1} = (\pi/2, \pi/2)$ and $q_{r_2} = (\pi/2, 3\pi/2)$. Further, $q^1_a(t) \in \mathbb{R}$ and $q^2_a(t) \in \mathbb{R}$ represents the agent's position at time $t$ along the two axes of the torus.

### Observation vector

The observation vector $u_c(t) \in \mathbb{R}^p$ used in Section 2.1.3 of the main text as control input to the dynamics of the control model is a concatenation of vectors $l_a(t) \in \mathbb{R}^6$, $x_r(t) \in \mathbb{R}^2$ and a scalar $h(t) = ||e(t)||_1 - ||u_p(t)||^2_2$, which represents the net amount of resources consumed by the agent at time $t$, such that $p = 9$. Firstly, $l_a(t)$ is given by

$$l_a(t) = [\sin^*(q_a(t)),\ \cos^*(q_a(t)),\ \dot{q}_a(t)]^\top \tag{17}$$

where $q_a(t) \in \mathbb{R}^2$ represent the position of the agent. The $\sin^*$ and $\cos^*$ functions represent an element-wise sine and cosine of a vector passed to them. Secondly, $x_r(t) \in \mathbb{R}^2$ from Section 2.1.2 of the main text represents the value of resources in the two patches. Thus we have

$$u_c(t) = [l_a(t),\, x_r(t),\, h(t)]^\top. \tag{18}$$

### System initialization

For each trajectory in a learning trial, the equations of the couple dynamical systems are initialized in the following way:

**Position model:** The initial values of $q_a$ and $\dot{q}_a$ in Section 2.1.1 of the main text are sampled from a random uniform distribution, distributed between $-\pi$ and $\pi$. Thus the agent can initially be spawned at any point on the surface of the torus with angular velocities of $\pm\pi$ rad/s along the two axes. The value of damping factor $\gamma$ is set to 0.3.

**Resource model:** The initial value of both the resources is set to 4.0. The value of the growth rate $\epsilon$ in Eq 5 of the main text is set to 0.1 by default. The value of decay rate $\sigma$ is always set to 0.01. Thus the carrying capacity which is defined as the maximum value to which a resource can grow becomes $\epsilon/\sigma = 10.0$. The consumption rate $\beta$ for the agent is set to 0.5.

**Control model:** The initial values of the membrane potential vector $x_c \in R^n$ as used in Eq 6 of the main text is set by multiplying the initial observation vector $u_c \in \mathbb{R}^p$ in Eq 18 to a matrix $R \in \mathbb{R}^{n\times p}$. Such that $x_c(0) = Ru_c(0)$. The number of neurons $n$ is set to 40 and, as highlighted in the previous section, the number of observations $p$ is set to nine. The values of all the learnable parameters $\theta$ are initialized using Glorot initialization [1]. In the case of the neuronal time constants as used in Eq 6 of the main text, we first initialized a value $c_i \in \mathbb{R}$ using Glorot-Normal initialization for the $i^{th}$ neuron. Then we used the equation $\tau_i = 10/(1 + e^{-c_i})$ to initialize the value of ${\tau_i}^{-1}$ which formed the elements of the $\alpha$ vector. This was done to maintain the value of time constants above zero for physical realism.

### Simulation details

The model was written using the Jax library, which utilizes just-in-time compilation coupled with an accelerated-linear-algebra compiler to implement computations in parallel [2]. The Diffrax [3] and Equinox [4] libraries were used to model the differential equations and solve them respectively. More specifically, the Tsit5 ODE solver [5] is used for numerical integration of the system equations. The solver further uses a proportional-integral-derivative controller with relative and absolute tolerances of $10^{-3}$ and $10^{-6}$ respectively, to control the step size of the simulations. The Optax library [6] was used for optimization during learning. We used an Adam optimizer to calculate the gradients and update the learnable parameters [7]. The learning rate (lr) for the optimizer was set to $3 \times 10^{-4}$ and the calculated gradients were clipped between the values of $-1.0$ and 1.0. All simulations were run on an Apple M1 CPU. A summary of all the hyper-parameters used for the learning procedure can be found in Table 1. Since all $n_b = 64$ trajectories are evaluated in parallel, following a vectorized approach, the computational complexity of the learning procedure per episode is proportional to the time taken by the solver to evaluate the system for a single trajectory. It depends heavily on the choice of the numerical solver and the stiffness of the system. In the case of opting for a forward Euler solver [8], the computational complexity scales similar to a discrete-time recurrent neural network [9].

**Table 1.** Hyper-parameters used for learning.

| Parameter | Value |
|---|---|
| Learning rate (lr) used to update $\theta$ | $3 \times 10^{-4}$ |
| Batch size | 64 |
| Initial step size | 0.04 s |
| Length of a trajectory | 1000 steps |
| Length of truncated interval for t-BPTT | 200 steps |
| Relative tolerance value for ODE solver | $10^{-4}$ |
| Absolute tolerance value for ODE solver | $10^{-6}$ |

### Code

The code used in this work can be found at `https://github.com/i-m-iron-man/NDE_Foraging.git`.
